# Robotic Laser Tissue Soldering for Damage-free Soft Tissue Fusion Guided by Fluorescent Nanothermometry

**DOI:** 10.1101/2024.02.29.582843

**Authors:** Oscar Cipolato, Tobias Leuthold, Marius Zäch, Georg Maennel, Sam Aegerter, Calinda Sciascia, Alexander Jessernig, Sima Sarcevic, Jachym Rosendorf, Vaclav Liska, Dennis Kundrat, Romain Quidant, Inge K. Herrmann

**Affiliations:** Nanoparticle Systems Engineering Laboratory, Institute of Energy and Process Engineering (IEPE), Department of Mechanical and Process Engineering (D-MAVT), ETH Zurich, Sonneggstrasse 3, 8092 Zurich, Switzerland; Particles Biology Interactions Laboratory, Department of Materials Meet Life, Swiss Federal Laboratories for Materials Science and Technology (Empa), Lerchenfeldstrasse 5, 9014 St. Gallen, Switzerland; The Ingenuity Lab, University Hospital Balgrist, Balgrist Campus, Forchstrasse 340, 8008 Zurich, Switzerland; University of Zurich, Faculty of Medicine, Rämistrasse 71, 8006 Zurich, Switzerland; Fraunhofer Research Institution for Individualized and Cell-Based Medical Engineering IMTE, 23562 Lübeck, Germany; Department of Surgery, Faculty of Medicine in Pilsen, Charles University, Alej Svobody 923/80, Pilsen, 32300 Czech Republic; Biomedical Center, Faculty of Medicine in Pilsen, Charles University, Alej Svobody 1655/76, Pilsen, 32300 Czech Republic; Nanophotonic Systems Laboratory, Institute of Energy and Process Engineering (IEPE), Department of Mechanical and Process Engineering (D-MAVT), ETH Zurich, Sonneggstrasse 3, 8092 Zurich, Switzerland

## Abstract

Minimally invasive surgical techniques, including endoscopic and robotic procedures, continue to revolutionize patient care, for their ability to minimize surgical trauma, thus promoting faster recovery and reduced hospital stays. Yet, the suturing of soft tissues ensuring damage-free tissue bonding during these procedures remains challenging due to missing haptics and the fulcrum effect. Laser tissue soldering has potential in overcoming these issues, offering damage-free seamless tissue fusion. To ensure the precision and safety of laser tissue soldering, we introduce feedback controlled fluorescent nanothermometry-guided laser tissue soldering using nanoparticle-protein solders within endoscopic and robotic contexts. Temperature-sensitive fluorescent nanoparticles embedded in the solder provide surgeons with immediate feedback on tissue temperatures during laser application, all while within the confines of minimally invasive (robotic) surgical setups. By integrating fluorescent nanothermometry-guided laser tissue surgery into endoscopic and robotic surgery, we pave the way for a new approach for safe and atraumatic soft tissue joining, especially in regions where traditional suturing is unfeasible.

## Introduction

Minimally invasive surgery (MIS), i.e., surgical procedures that use various techniques to minimize the size of the needed incisions, has been growing in popularity for many years, with case increases of up to 462 % between 2000 and 2018.(*1*) However, due to the limited working space inside the patient associated with this surgical technique, the disadvantages of suturing become more pronounced.(*2*) Difficulties in suturing in minimally invasive surgery are greatly amplified by the fulcrum effect (the result of pivoting instruments at varying depths), minimal or even absence of haptic feedback, amplified tremor and a reduced surgical field view, among other factors.(*3*, *4*) To overcome some of the limitations in conventional MIS, robotic MIS has been introduced to the operating rooms with great success in the last decades.(*5*, *6*) However, sutures and staples have their own set of intrinsic drawbacks, particularly in soft tissue injuries to the liver, the intestine, blood vessels, nerves or the dura mater. Suturing and stapling techniques are not only susceptible to inflammation, infection, and slow healing, but they can also cause tissue trauma(*7*) and lead to dangerous fluid leaks.(*8–10*)

To overcome the current limitations of suturing in endoscopic and robotic surgery, various surgical glues and wet tissue adhesives have been investigated.(*11*, *12*) While potentially promising, limitations(*13*, *14*) include toxicity (e.g. cyanoacrylates(*15*)), immunogenicity, low adhesion (e.g. fibrin- or poly(ethylene glycol)-based glues(*16*)), and swelling (e.g. hydrogel-based sealants(*11*, *12*, *17*)). Most of these technologies have found their main application in open surgery and they are poorly applicable for minimally invasive procedures. The physical dimensions and application techniques of these adhesives frequently pose challenges for seamless adaptation to a minimally invasive environment.

Laser tissue soldering represents a promising alternative: it is a surgical technique that combines laser light with a solder paste, a temperature-activatable adhesive material, usually protein-based, to create strong watertight bonds.(*18–20*) The attractiveness of soldering lies in its ability to rapidly form bonds with good strength,(*21–23*) low inflammatory response,(*24*) reduced scar tissue formation,(*25*) and reduced access to pathogens thanks to the creation of a waterproof seal.(*26*) Furthermore, this technique is, in principle, readily adaptable for minimally invasive procedures.(*10*) The minimal amount of material required for bonding, the straightforward guiding of laser light to the bonding location, and the simplicity of the technique — which does not require intricate motions or exhaustive training — all contribute to its feasibility.(*27*, *28*) Although laser tissue soldering holds great promise, it remains experimental.(*19*) The technique’s success hinges crucially on the precise temperature management during the procedure. For optimal bonding, temperatures between 60 °C and 80 °C are required, yet the surrounding tissue must remain significantly cooler to avoid irreversible tissue damage. Insufficient temperatures during soldering result in inadequate protein-tissue bonding, while excessive heat can damage surrounding tissues and delay healing.(*18*) Thus, mastering the confinement and control of the temperature is critical for clinical translation.

In this work, we introduce a new minimally invasive wound closure technique for (robotic) MIS, based on controlled laser tissue soldering guided by fluorescent nanothermometers able to create seamless water-tight sealing. Nanothermometers have recently been successfully integrated into proteinaceous solders,(*29*) and offer non-invasive temperature measurements. The proteinaceous solder material can further be supplemented with nanoabsorbers, including gold nanorods or titanium nitride nanoparticles(*29*) for optimal differential heating. As a critical step towards more accurate and less invasive soldering, we demonstrate automatic temperature control and smart laser power modulation for high-performance laser tissue soldering in (robotic) MIS, enabled by an intelligent solder paste that contains NIR-fluorescent nanothermometers and nanoabsorbers. Moreover, we present integrating computer-vision based solder recognition along with multiple approaches to guide the surgery by measuring the nanothermometry-based temperature distribution during soldering for optimal outcome. Finally, we demonstrate the crucial advantages of minimally invasive soldering in clinically-relevant scenarios.

## Results and Discussion

### Design of a controlled soldering process for minimally invasive procedures

To address the current limitations of soft tissue joining in MIS, the ideal approach should not only facilitate wound closure in a minimally invasive manner but also establish robust and reliable tissue bonds, all while preserving the integrity of the surrounding healthy tissue. Good reliability can be obtained only if the technique is easy to perform and relies on a robust environment-independent mechanism. Moreover, it should be compatible with existing surgical tools of widespread use (Figure 1a, b). Amongst the potential technologies for soft tissue repair in the constrained environment of MIS, laser tissue soldering offers a particularly promising approach. Especially using proteinaceous solders with integrated nanoabsorbers for optimal heat confinement, and nanothermometers for accurate temperature control during soldering, offers a route to seamless tissue fusion.

**Figure 1:**
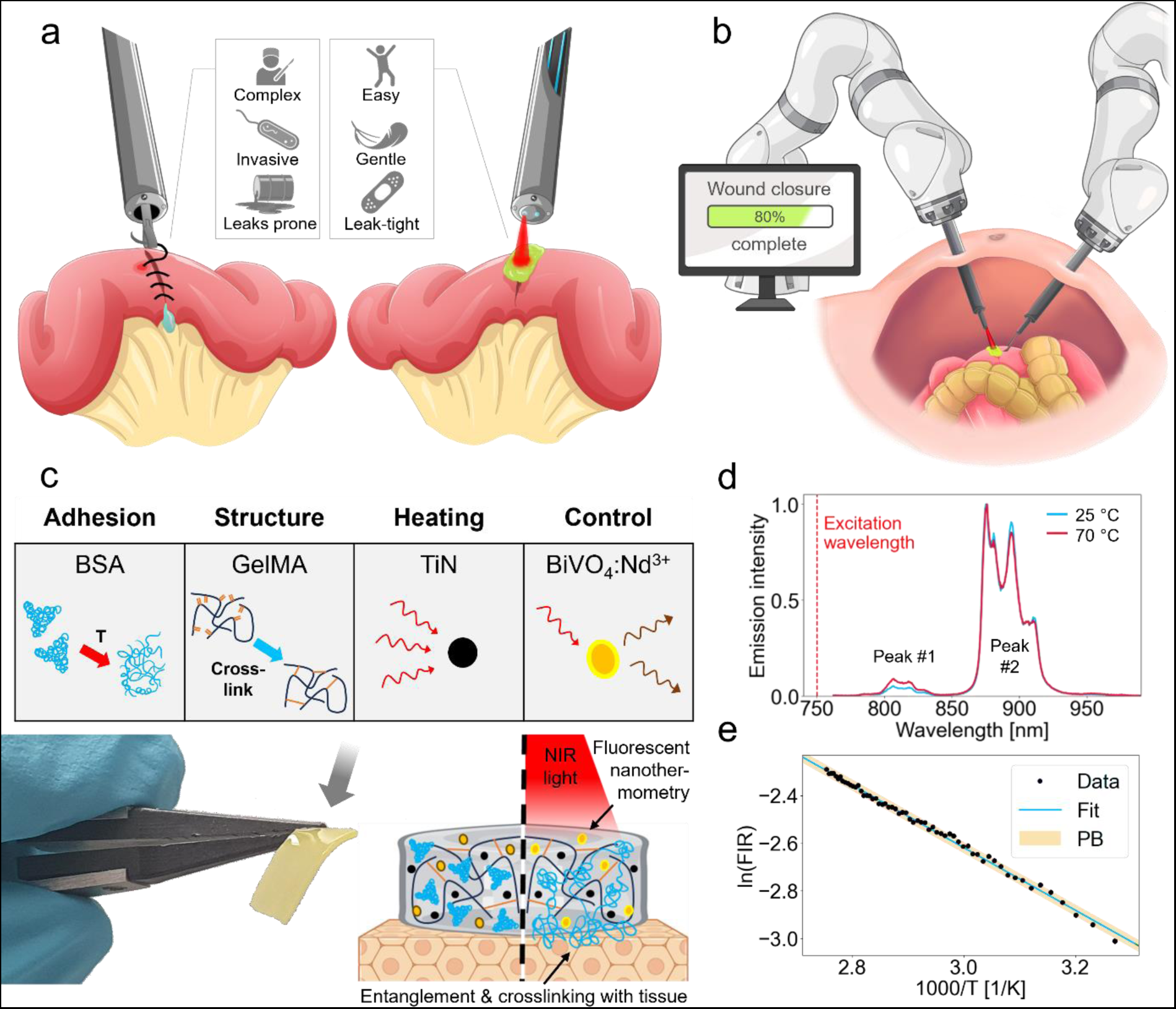
Controlled tissue soldering for seamless tissue fusion in MIS. (a) Illustration showing the advantages of laser tissue soldering compared to conventional suturing of soft tissues. (b) Schematic of laser tissue solder integration into robotic minimally invasive surgery. (c) The components of the solder paste (shown in the photo) are illustrated schematically, along with a schematic of the soldering mechanism. (d) Fluorescence spectra collected at two different nanothermometer temperatures (normalized intensity). The ratio of the two fluorescence emission peaks of BiVO_4_:Nd^3+^ are used for non-contact temperature measurements. (e) Calibration plot used to calculate the temperature from the fluorescence intensity ratio FIR (a = 1.33 ± 0.03, b = -1.32 ± 0.01 K^-1^ are the values of the fit *y* = *a* + *b* × *x* with *y* = ln(*FIR*) and *x* = 1/*T*; PB = Prediction Band).

We designed an albumin-based solder paste optimized for MIS, by using gelatin-methacrylate (GelMA) as a structure-giving moiety for optimal endoscopic and robotic application and manipulation of the paste. Addition of GelMA as a structure-giving element is essential to avoid uncontrolled liquefaction of the solder material during manipulation. Additionally, TiN (for confined temperature increase) and Nd:BiVO nanoparticles (for nanothermometry) were integrated into the paste for optimal performance and integration into feedback controlled laser tissue soldering (Figure 1c). TiN nanoparticles are used as nanoheaters due to their broad absorption spectrum, high photothermal efficiency, stability, and low cost. Flame-made bismuth vanadate doped with neodymium (BiVO_4_:Nd^3+^) serve as fluorescent nanothermometers. Their merit lies in their excitation and emission wavelengths situated within the near-infrared (NIR) biological window, ensuring safety and robust performance regardless of the environment (Figure 1d). The fluorescence intensity ratio (FIR) can be calculated from two BiVO_4_:Nd^3+^ emission peaks and correlated with the temperature through the Boltzmann thermal equilibrium equation (Figure 1e and Materials and Methods). Using a ratiometric approach brings clear advantages, as it makes the measurement unaffected by nanothermometer concentration, laser power, and weakly-absorbing materials in the NIR situated between or on the paste and the measuring fiber, such as thin tissues or various bodily fluids. This is an important distinction from widely used thermometry methods that rely on intensity measurements in the Mid-IR range (3-20 µm, and most commonly 3-5 µm or 8-14 µm),(*10*) which are strongly affected by environmental factors. The absorption coefficient of water in this range is in the order of 10^4^-10^6^ m^-1^(*30*), about 4 orders of magnitude higher than in the NIR range (absorption coefficient of water between 0.7-1.4 µm is in the order of 10^0^-10^2^ m^-1^). This means that with infrared thermometry a water layer of 1-100 µm on the paste or on the tip, something that readily occurs during MIS due to tissue manipulation, would mask the actual temperature, leading to incorrect measurements for conventional thermometry, while fluorescence nanothermometry measurements remain unaffected.

Soldering using the nanomaterial-enhanced solder materials can readily be integrated into a (robotic or endoscopic) MIS setup. By integrating two optical fibers in an endoscopic instrument, the solder paste can be irradiated through one fiber and the fluorescence of the nanothermometers can be measured simultaneously by the second fiber, providing crucial information about the temperature during soldering in a contact-free, minimally invasive manner (Figure 1b). Filtering of the light source and of the signal can be performed at the distal end of the optical fibers, away from the soldering location, allowing for increased flexibility on how to filter and process the signal collected from the surgical site within the patient. For example, the temperature information can be used to modulate the laser power to provide controlled and safe soldering, either in an automatized or user-controlled fashion. Moreover, the optical fibers can be removed and replaced or sterilized, a crucial requirement for in vivo studies and use in clinical operating rooms.

### Automatized feedback for controlled heating

Aiming to attain increased precision and reliability in laser tissue soldering, the adoption of an automatic feedback control system emerges as an appealing choice. This preference arises from the capacity to eliminate user-dependent variables, thereby improving the reliability and safety of the process. Thus, we implemented a Proportional-Integral-Derivative (PID) controller to maintain the temperature at the target site within the patient’s body by regulating the laser power, driven by real-time temperature measurements provided by fluorescent nanothermometry (Figure 2). PID controllers are an established feedback control loop used in industrial applications, thanks to their ease of integration and built-in stability. Their efficacy is further underscored by the tunability of three key parameters: proportional gain (P), integral gain (I), and derivative gain (D). These parameters allow for customization of controller responsiveness and speed, facilitating optimization for speed or the mitigation of overshooting, contingent on specific procedural needs. By optimizing the parameters, the controller can reliably maintain target temperatures in the range relevant to laser tissue soldering while minimizing rise times (below 10 s) and showing no overshooting (Figure 2a). For applications in MIS, performance under a variety of different application scenarios is essential for patient safety and outcome. Appropriate tailoring of the controller gives resilience to the system to vertical movements, reaching and maintaining the target temperature, albeit with variations in rise time when distances undergo significant changes (Figure 2b). The parameters of the controller can further be tuned to make it more responsive, reaching the target temperature much faster and showing only a small overshoot (Figure 2c).

**Figure 2:**
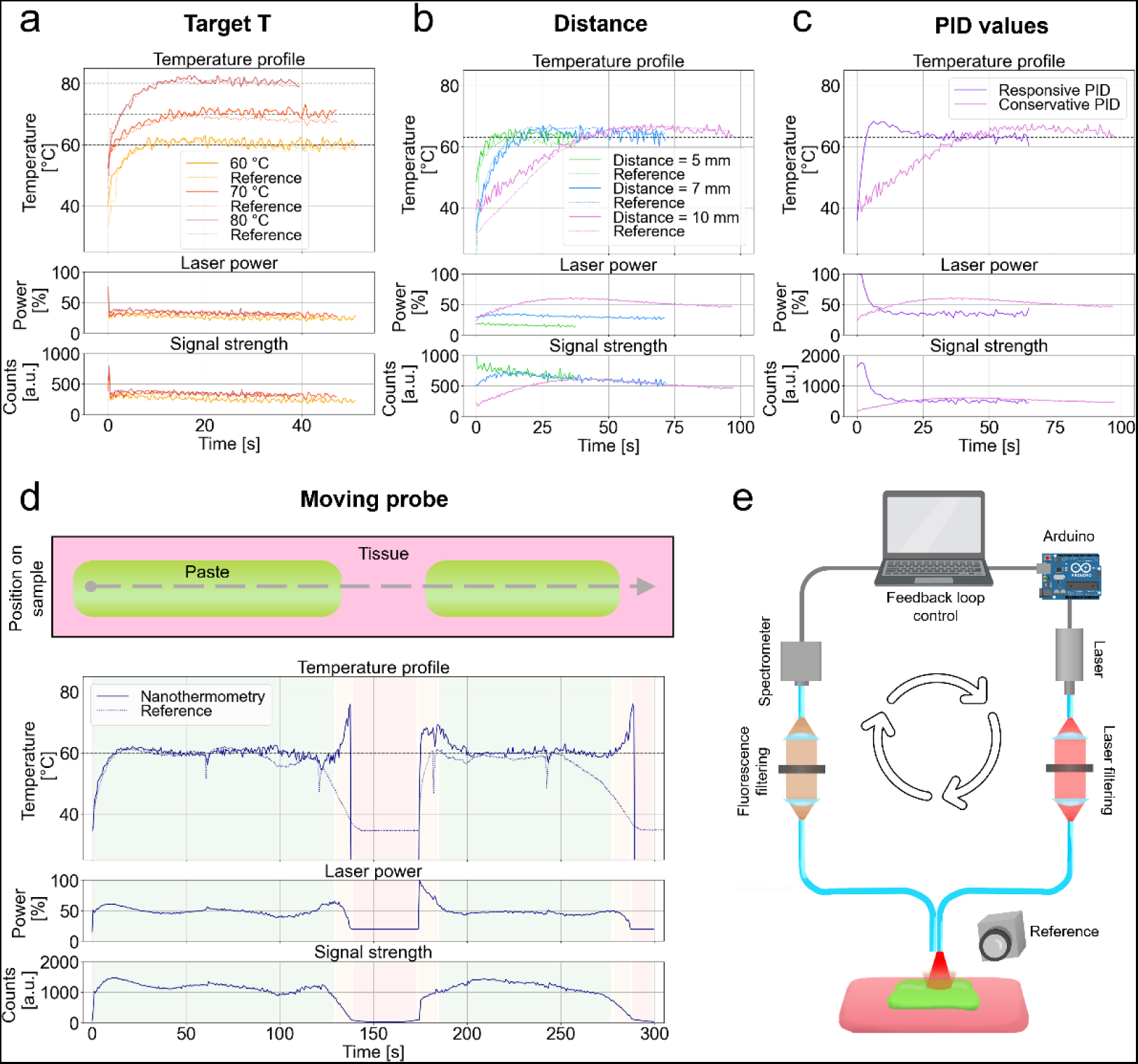
Automatized feedback for controlled local heating. (a) Feedback-controlled soldering allows for soldering at various temperatures (60, 70, 80 °C), achieving the target temperature in less than 10 s for temperatures needed in tissue soldering (top plot). Temperature control is kept by modulating the laser power (middle plot) and based on the signal strength (bottom plot). (d = 10 mm) (b) The target temperature (here 63 °C) is reached also at different distances of the fibers from the solder paste, however, with a slower response at higher distances. (c) The PID parameters can be tuned to obtain a responsive behavior, at the cost of an initial small temperature overshoot. (d = 10 mm, T = 63 °C) (d) The feedback mechanism can be employed also while the optical fiber-endoscope is moved over the wound (as illustrated in the top part): two pieces of solder paste are placed on a piece of pig intestine and the fibers are moved over them. The controller is also able to discriminate when the paste is being irradiated (green areas) and when it is above healthy tissue (red areas), automatically lowering the laser power to a safe level (15%). (d = 10 mm, T = 60 °C). (e) Illustration of the feedback loop setup used for paste recognition and soldering: the solder paste is placed on ex vivo tissue (pig dura mater), one fiber is used for laser excitation, one fiber is used for fluorescence measurement, and a thermal camera is used as reference. The nanothermometry spectra measured by the spectrometer are used to control the feedback loop through an Arduino that modulates the laser power.

Furthermore, we demonstrate that this technology succeeds when navigating diverse sections and complex topologies of a wound (Figure 2d,e). For demonstration, two pieces of solder paste 2 mm apart are placed on a piece of tissue, while the laser is moved at a constant speed (2 mm/min) across them. The soldering temperature is maintained at the set target temperature of 60 °C throughout the entire section on the solder paste (green-shaded area in Figure 2d). Upon detection of healthy tissue and areas devoid of solder material, the controller adjusts laser power to a secure level, averting potential damage, as opposed to implementing an exponential increase. Such a feature is possible thanks to the fluorescence emission signal emitted by the paste. When the signal strength is low, the laser power is switched to a safe level that still allows fluorescence detection (red-shaded areas in Figure 2d), which ensures that the soldering restarts once on the correct location (green-shaded areas in Figure 2d). Moreover, by measuring the reflected excitation light, one can distinguish between being too far from the paste and being too close to tissue. At the edges of the paste (yellow-shaded areas in Figure 2d) the temperatures tend to be overestimated due to lower signal-to-noise ratio; however, this does not represent a problem, as the laser power gets lowered, rendering the process safe.

### Machine learning for low-cost thermal distribution imaging

During soldering, the temperature distribution provides valuable information for estimating the size of the heat-affected area and how quickly previously heated areas are cooling down. Fluorescent nanothermometry requires detailed spectral information. However, a simple setup using only a fiber-coupled laser and spectrometer lacks sensitivity to spatial differences. Temperature distribution information can be used to ensure a more uniform soldering throughout the entirety of the solder paste, as well as avoiding local hot spots that can damage the tissue. Spatially resolved 2D spectra can only be captured with specialized cameras like hyperspectral ones, which are relatively expensive. This applies to nanothermometry based on ratiometric methods with closely spaced peaks, as well as approaches involving peak shifts and fluorescence decay time.

In this context, we introduce a machine learning approach based on Convolutional Neural Networks (CNN) combined with a lens-free, low-resolution fiber bundle probe (Figure 3) for thermal distribution measurement in MIS. Using a fiber bundle with a small number of fibers, such as a 5×5 square array, results in a low-resolution hyperspectral image. These low-resolution images can be challenging to interpret and do not offer significant advantages when presented to professionals performing the procedure. Furthermore, they tend to overestimate the temperature at the edges of the soldered area, primarily because the signal primarily originates from illuminated portions of the paste, which typically have higher temperatures. Our developed upscaling CNN, based on emerging architectures (*31*), can accurately reconstruct the thermal distribution and increase the resolution by a factor of 4. This substantial resolution improvement is achievable due to the limited diversity of temperature distribution profiles during laser irradiation. To train the CNN, we first generated a dataset by simulating various temperature distributions and their corresponding fluorescence signals detected by each fiber tip, including simulated noise. While Structural Similarity Index (SSIM) loss and peak signal-to-noise ratio tend to improve with increasing epochs, the validation loss suggests that the optimal training performance occurs at around 250-300 epochs. Therefore, it is advisable to select this range to avoid overfitting to the training data (Figure 3c). The CNN demonstrates strong performance in reconstructing the thermal distribution’s position, size, and range (including peak temperature) within the training dataset, even when the input data contains limited information (Figure 3d). We further validate the CNN’s performance using experimental data. A stage is employed to position a fiber tip where each fiber in a given bundle (in this case, a 5×5 square array of fibers side-by-side) would be located, allowing us to collect light at various positions. The fluorescence signal is then used to reconstruct the thermal distribution, which is compared to a reference thermal distribution obtained using a thermal camera (Figure 3f). Importantly, the thermal distributions calculated using the CNN match the measured distributions very well and the peak temperatures are predicted with high accuracy (*ΔT_max_* = 2.6 ± 1.0 °C, SSIM = 0.85 ± 0.04, *N* = 3). This approach is also directly applicable to fluorescent nanothermometry based on peak shifting or lifetime measurements, where precise spectral information is required and provides a low-cost route to thermal distribution imaging in MIS.

**Figure 3:**
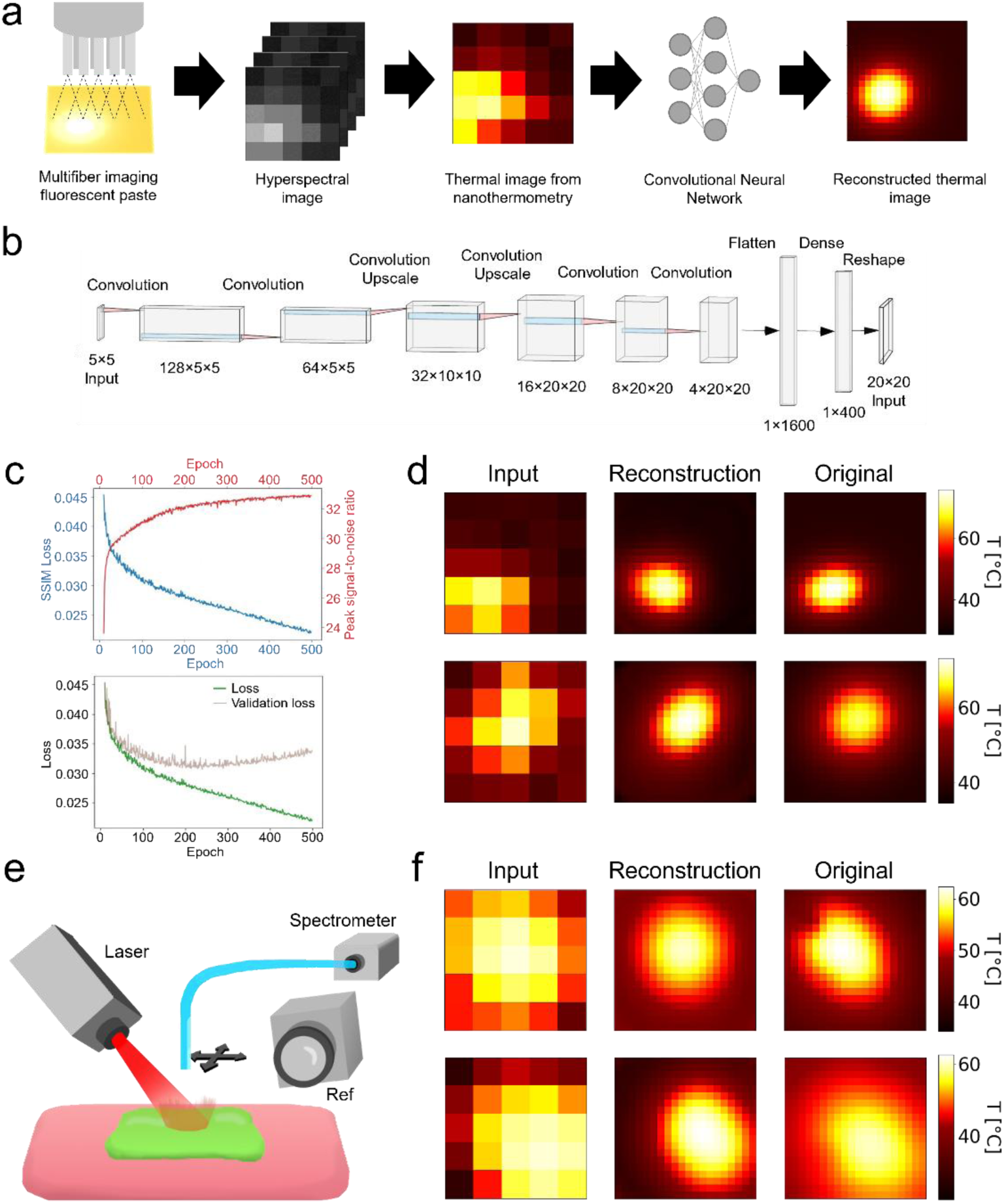
Machine learning for low-cost thermal distribution imaging. (a) Illustration of how a multifibre system coupled with the upscaling Convolutional Neural Network (CNN) can reconstruct the thermal distribution during laser excitation starting from a low-resolution hyperspectral image. (b) Upscaling CNN architecture. (c) Performance of the trained CNN showed with training loss. (d) Examples of simulated data reconstructed using the upscaling CNN. (e) Experimental setup used to collect the fluorescence from each individual fiber featuring inexpensive equipment: the fluorescence light is collected by a displaceable fiber and is lead, after being filtered, to a spectrometer. The signal is used to calculate the temperature which is compared to a reference thermal camera. (f) Examples of experimental data reconstructed using the upscaling CNN. (FOV = 0.25 cm).

### Thermal distribution imaging with image-guiding fiber and NIR cameras in robotic soldering

Alternatively, instead of using the upscaling algorithm and spectrometer setup, image-guiding fibers (also known as leached fibers) can be employed in combination with NIR cameras to obtain valuable thermal distribution information in a minimally invasive manner (Figure 4). Image-guiding fibers are thin bundles of thousands of fibers arranged orderly, allowing them to transmit images from one side of the fiber to the other with minimal signal loss and cross-talk between fibers.(*32*) To retrieve the temperature distribution, we use a dichroic beam splitter to separate the two emission peaks and two NIR cameras. A GRIN lens (gradient-index lens) is used to create an image on the object side, while a microscope objective captures images on the distal side of the fiber with the two cameras (Figure 4a).

**Figure 4:**
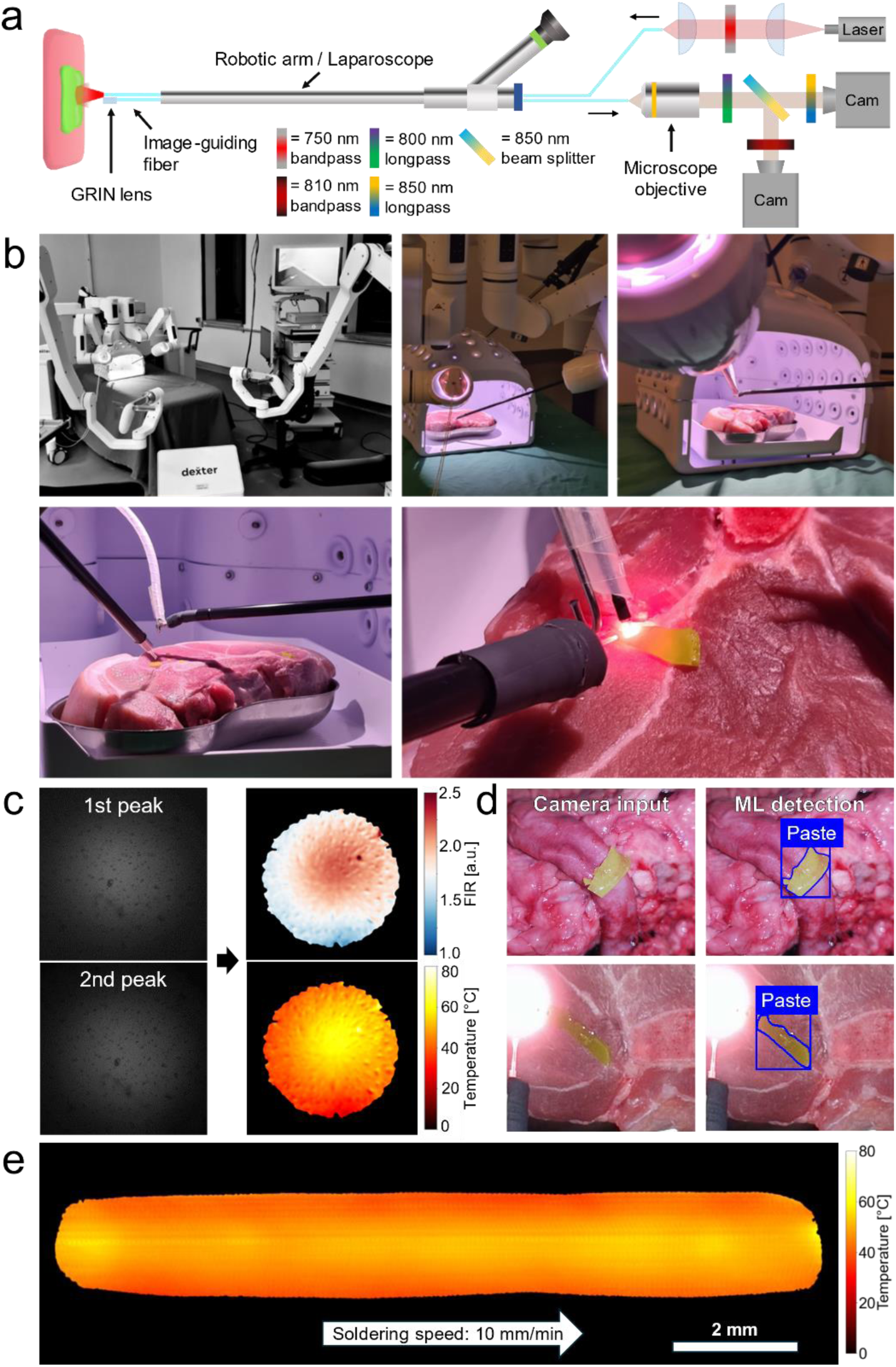
Multicore fiber thermal imaging. (a) Schematic of the setup used to measure the thermal distribution during soldering, featuring two NIR cameras to detect the fluorescence of the two emission peaks. (b) Pictures of the robotic system (Dexter, Distalmotion SA) with integrated tissue soldering equipment. The dual-fiber system for automatized feedback-controlled soldering is incorporated with the instrument of the left robotic arm, while the imaging fiber is spatially manipulated using a gripper instrument of the right robotic arm. (c) Example of images recorded with the two different filters during soldering of a piece of solder paste. The FIR is calculated from the two images that represent the fluorescence of the two peaks (acq. time = 500 ms). The FIR is then used to compute the temperature distribution of the solder paste (FOV = 0.6 cm). (d) A machine learning algorithm is developed and applied to automatic detection and localization of the paste during surgery, information that can be used to automatically define and generate a soldering path. Top: image of the paste on a porcine aorta used for training. Bottom: picture taken by a laparoscope at the start of robotic soldering. (e) The thermal imaging through fluorescence can be implemented together with the automatized feedback-controlled soldering. This reconstructed image, made by adding images taken while the soldering probe was moving, shows how the temperature is kept at the target temperature (60 °C) throughout the entire soldering path.

By adjusting the position of the GRIN lens relative to the fiber tip, we can change the focus and field of view (FOV). The choice of BiVO nanothermometer is particularly well-suited to this method because its two thermally coupled emission peaks are spectrally distant (more than 40 nm apart) and can be clearly separated using commercially available dichroic beam splitters, bandpass, and longpass filters. Additionally, the used wavelengths fall within the sensitivity range of CCD and CMOS cameras, making it possible to utilize established and cost-effective technology, as opposed to InGaAs or Mid-IR image sensors.

This system can also be coupled with the automated feedback for temperature control and be readily applied to robotic surgery (Figure 4b). The flexible and thin fibers can be easily incorporated with the instrument of a surgical robot (here exemplarily shown using Dexter, Distalmotion SA), and their position can be easily controlled through the remote robotic interface. The surgical workflow is as follows: First, the paste is deployed through one of the trocars and placed on the area of interest using the articulated robotic gripper, then, soldering is started by moving in place the instrument with the two fibers for the feedback-controlled soldering. The image-guiding fiber is inserted through another trocar and moved to the soldering site for monitoring purposes by the gripper arm of the surgical robot. During soldering, the thermal distribution can be seen thanks to the image-guiding fibers. The two NIR cameras capture fluorescence images with the same acquisition time (0.5 s) but different gain values, given that the first peak (at 820 nm) is dimmer than the second peak (at 870 nm). Based on the material intrinsic calibration curve and considering the different gain values, the FIR distribution can be converted into the thermal distribution of the paste (Figure 4c). Only temperature measurements from areas with sufficient signal strength are displayed.

Information about the spatial paste position can be given to the robotic system to enable soldering in an autonomous fashion, e.g., generating temperature-optimal trajectories. To this aim, a machine learning image segmentation algorithm compatible with real-time applications was developed. The solder paste detection was achieved by using this algorithm, able to reliably recognize and locate the solder paste even in images taken by the robotic laparoscope camera (Figure 4d, Supplementary Information). Such information can be used to automatically define a soldering path with the aim of soldering an extended region, such as the region along a cut or a suture line. Moreover, thanks to the automatic temperature control, the temperature can be kept at the desired value even while the soldering setup is moved along the soldering path in a fully automated manner.

This is demonstrated exemplarily by the temperature map acquired by the multifiber imaging system (Figure 4e), where the various temperature distributions acquired during soldering of a 13 mm-long piece of solder paste are presented. Each frame can be linked with its location on the paste, allowing the creation of the temperature map of the paste during soldering. Such a map can be used to ensure optimal heating of the entire target area, avoiding local overheating spots or areas with suboptimal heating that would hinder proper tissue bonding, and can be fully integrated into the robotic setting. For example, robot control can be complemented by a cognitive supervisory framework(*33*) to increase the level of task autonomy and facilitate uniform area heating at targeted temperatures realizing homogeneous tissue bonding.

### Minimally invasive laser tissue soldering in vivo

The potential of iSoldering for MIS is further demonstrated in an in vivo porcine model with torso dimensions approximating the ones of human patients (Figure 5). The optical fibers for soldering can be easily incorporated into a typical laparoscopic setup by inserting them into a 5 mm trocar (Figure 5a). To insert the solder paste inside the abdomen, the solder paste can be placed in the groove of a grasper, without the need for other specialized equipment (Figure 5b). With this modality, the paste is protected by the grasper during insertion, allowing it to maintain its integrity throughout the insertion process even through narrow trocars. The paste is then taken using a second grasper and then positioned where needed, such as on a piece of intestine (Figure 5c). The intestine was chose as it is one of the most challenging organs for laparoscopy. The use of crosslinked GelMA instead of normal gelatine is of key importance in laparoscopy, as the working environment is kept at body temperature, causing the melting of non-crosslinked gelatine-based paste in less than a minute. By using crosslinked GelMA the paste does not melt and can be easily placed on the wound by the graspers. The optical fibers can be directed towards the area to solder by moving the trocar through which they are inserted and sliding them in or out to adjust the fiber-tissue distance. Feedback-controlled soldering can be safely initiated when the fiber is approximately in place. The algorithm will ensure that high laser powers are used only when the solder paste is detected (Figure 5c). When on the surrounding healthy tissue, the laser power will be reduced to avoid damage, but its presence will provide useful information to guide the fiber positioning. The optical fibers can also be moved with the help of a grasper. By directing the fibers with a grasper, areas that are more difficult to reach can be illuminated and soldered, such as damaged areas on the lateral abdominal wall (see SI). The effectiveness of feedback-controlled iSoldering in MIS is further demonstrated based on histological data (Figure 5d). By comparing the soldered tissue with the healthy tissue control, no visible damage is observed on the soldered tissue and a seamless and firm bonding of the solder paste to the tissue is observed. Good adhesion is further demonstrated based tensile strength testing performed on porcine intestine, achieving around 40% of the strength of intact tissue (Figure 5e).

**Figure 5:**
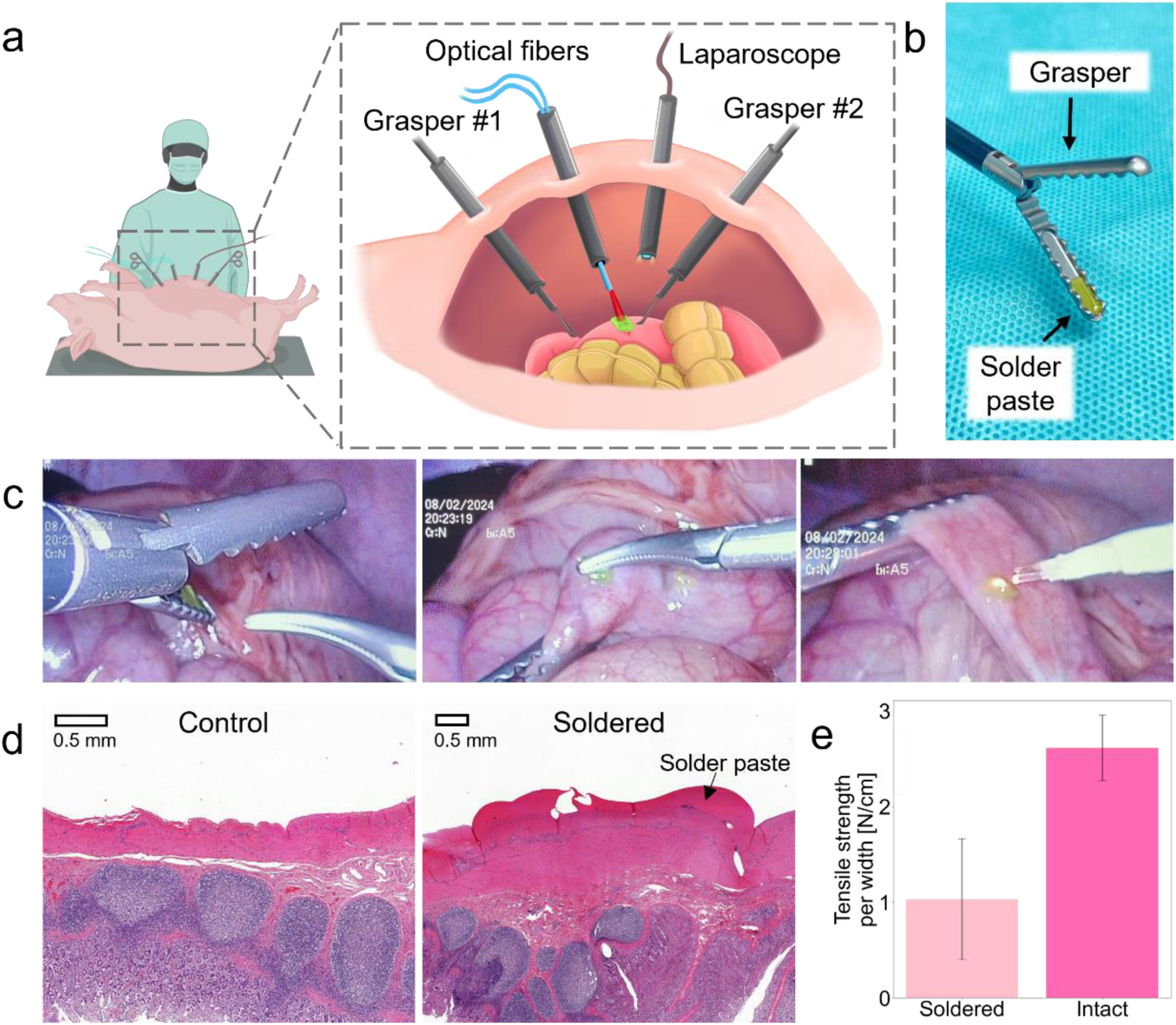
Application in laparoscopic surgery in vivo. (a) Soldering performed laparoscopically on an in vivo porcine model, with the optical fibers passing through a trocar. (b) The solder paste is placed in the groove of a grasper for easy insertion inside the abdominal cavity. (c) Then, the paste can be taken and positioned with a second grasper. The optical fibers are directed to the paste and soldering is performed. (d) Histology of the soldered intestine compared with a healthy control, showing no visible thermal damage and good adhesion. (e) Tensile strength values of soldered intestine compared to intact intestine.

## Conclusions

This work presents the integration of nanothermometry-guided laser tissue soldering into the minimally invasive surgical procedures, addressing significant surgical limitations that have persisted hitherto. The inherent delicacy and complex structure of soft tissues often render conventional mechanical fusion techniques like suturing and stapling inadequate or impractical. The presented approach of laser tissue soldering, guided by real-time fluorescent nanothermometry, emerges as a compelling solution to these challenges, opening new avenues for secure, precise, and controlled soft tissue fusion in MIS, including robotics. By leveraging the precision and real-time feedback offered by fluorescent nanothermometry, laser tissue soldering overcomes traditional barriers, presenting a straightforward, reliable, and adaptable methodology conducive to seamless incorporation within laparoscopic, endoscopic, and robotic surgical systems. This intricate thermal control, and its integration into robotic navigation mitigates the risks associated with overheating or underheating, thus substantially augmenting the safety and usability of the procedure. Additionally, the minimally invasive thermal imaging capability augments the visualization of the soldering process, offering surgeons an enhanced perspective and a higher degree of control, making this technology not only safe but highly user-friendly. Moreover, the developed platform could potentially benefit the field of hyperthermia at large (for cancer and other pathologies), where precise light dosing is also crucial.

The research presented in this paper paves the way towards safe damage-free automatized surgery on fragile internal tissues. After many years of research on laser-based tissue bonding, the proposed approach is slowly breaking the barrier that is preventing this technique from becoming widely accepted in clinical practice. Further investigation and development on data-driven automation will allow for unprecedented wound closure reliability and reproducibility, which will have to be tested on various organs and validated in surviving animal experiments.

## EXPERIMENTAL

### Paste formulation

The solder paste was adapted based on the formulation reported in (24), however, GelMA was added for improved properties during robotic manipulation and reduced fluidity at body temperature and during the soldering process. Unless specified, all concentration percentages are expressed as mass of solute over mass of solvent (water). Briefly, water solutions of bovine serum albumin (BSA, Sigma– Aldrich, A2153), gelatin (from porcine skin, gel strength 300, Type A – Sigma–Aldrich, G2500), flame-made BiVO_4_:Nd^3+^ nanoparticles, and TiN nanoparticles (PlasmaChem GmbH, PL-HK-TiN) were mixed and then casted into the appropriate mold. The paste was cooled for at least 1 h at 4°C and then kept at that temperature until use, not later than 2 weeks after production. The pastes that contained GelMA were produced using gelatin modified with methacryloyl (GelMA) instead of normal gelatin. The GelMA was synthetized using 0.0069% LAP (Lithium phenyl-2,4,6-trimethylbenzoylphosphinate, Sigma–Aldrich, 900889) was used as crosslinking initiator and added to the solution as last component. The paste was first left for at least 1 h at 4 °C and then crosslinked under UV irradiation for 5 min (UVASPOT 400T, Dr. Hönle AG). Crosslinking of the paste prevents melting before BSA denaturation.

### Nanoparticle characterization

The calibration of BiVO nanoparticles was carried out calculating the fluorescence intensity ratio of the two fluorescence peaks. The two peaks were defined as the regions between 790 – 840 nm and 840 – 945 nm respectively. Transmission electron microscopy (TEM) of BiVO and TiN nanoparticles was carried out using a transmission electron microscope (EM900, Carl Zeiss Microscopy GmbH) at 80 kV. Holey carbon-coated copper grids (200 mesh, EM Resolutions) were incubated with poly-l -lysine solution (P8920, Sigma–Aldrich) for 10 min and subsequently washed with ultrapure (milliQ) water. The BiVO and TiN nanoparticles samples were first dispersed in milliQ water and subsequently drop-casted onto the grids.

### Feedback loop: optical setup

As excitation and heating source, a 750 nm CW laser (MDL-III-750-2W, CNI Lasers) with a maximum output power of 2 W was used. The power of the laser was controlled with pulse-width-modulation (PWM) via an Arduino microcontroller (Arduino UNO). A sturdy portable optical module was used to increase the monochromacity of the laser light. This module includes: an optical fiber (NA 0.22, ⌀400 µm) to guide the laser from the laser head to the optical module; a collimating plano-convex lens (f = 30 mm, 780 nm V-coating; Thorlabs); a 750 nm bandpass filter (FWHM = 40 nm; FB750-40, Thorlabs); a plano-convex lens (f = 30 mm, 780 nm V-coating; Thorlabs) for coupling with the output fiber. A medical optical fiber (NA 0.37, ⌀600 µm, 1 m length; Wuhan Medfibers Technology Co.) was used to deliver the laser light to the target. After passing the optical module and the medical fiber the maximum available power was reduced to 1.22 W.

To record the fluorescence a NIR spectrometer (100 µm slit; STS-NIR, Ocean Optics) was used. A medical optical fiber (NA 0.37, ⌀600 µm, 1 m length; Wuhan Medfibers Technology Co.) was used to collect the fluorescence light from the target. The medical fiber was connected to a sturdy optical module used to filter the fluorescence light. This module includes: a collimating achromatic doublet lens (f = 30 mm; Thorlabs); a 785 nm longpass filter (47-508, Edmund Optics); an achromatic doublet lens (f = 30 mm; Thorlabs) for coupling with the fiber (NA 0.22, ⌀550 µm) connected to the spectrometer. A thermal camera (PI 640i, Optris) was used to record reference data. Additionally, the tips of the fibers were fed through the aperture of a standard 5 mm trocar, to showcase the compatibility with common laparoscopic instruments. Apart from when otherwise stated, the same optical components (laser, spectrometer, filter and fibers) were also used in the other experiments.

### Feedback loop: controller

The controller algorithm was build based on the Python module “simple-pid v2.0.0” by Martin Lundberg.(*34*) A minimum laser power is set between 10-20%, depending on the configuration used, which is also used a starting laser power. The acquisition time is set between 300 and 800 ms.

To derive the calibration a data set consisting of fluorescence spectra and their corresponding temperatures is needed. The calibration can be performed either using a heating plate or using the laser itself as source of heat. Individual spectra were acquired in regular intervals and the reference data was taken with a thermal camera. The calibration was repeated with each paste sample for increased accuracy.

The general workflow is here presented:

1. The program measures a spectrum to correct for any background signal that is present. This step is repeated after a user-defined number of iterations (default: 100 iterations).
2. The laser is started with a pre-specified minimum power (default: 20 %). The minimum power is chosen depending on the working distance and heating characteristics of the paste. It can be reduced when the temperature keeps on rising at minimum power, while it may be raised to increase fluorescence signal, such as at long working distances.
3. The spectrum is taken, with a user-defined integration time (default: 500 ms). The integration time may be adjusted based on signal strength, with lower values yielding better controller response at the cost of SNR. The integration time is the main limitation of the loop sample rate.
4. If the number of counts is too low (default: 150 peak counts), the laser is switched to the minimum power to avoid heating healthy tissue.
5. The temperature is calculated and fed into the PID regulator, returning a new value for the laser power. The PID gain parameters can be adjusted during run time, with presets available.

The PID controller was tuned heuristically, with the following main performance requirements, in order of importance: stability in all scenarios; overshoot lower than 5 °C; *T*_90_ (time to rise to 90% of the set value) lower than 20 s. Setpoint-tracking was deemed secondary since the target temperature will most likely be held constant in LTS operation, putting the focus instead on disturbance rejection (movement during soldering).

### Feedback loop: soldering

The performance of the system was evaluated in different scenarios. In all cases, a solder paste with 25% BSA, 6.5% Gelatin, 0.3% BiVO and 0.002% TiN was used. The solder pastes were placed on top of a piece of ex-vivo porcine dura mater (retrieved from the local slaughterhouse), which was placed on a moving stage (a modified 3D printer). The distance of the sample from the fiber tip was changed between 0.5 and 1.5 cm, while the target temperature was changed between 50 and 80 °C. In the case of the moving soldering, the sample was moved at the speed of 2 mm/min.

### CNN training dataset creation

The process used to generate simulated data to train the CNN for the reconstruction of temperature distributions in a luminescence nanothermometry temperature profile is here explained. The training data must have the measured thermal images (i.e., 5×5 images) and the respective original 640×480 thermal images. A simulation algorithm that simulates the fluorescence received by an optical fiber placed in front of a laser-excited surface with BiVO nanoparticles was developed and used. The simulation algorithm for data creation is divided into three parts: database creation, modelling, image formation.

In a first step the algorithm is building a temperature-spectrum database. Using a thermally controlled cell (qpod 2e, Quantum Northwest), a 0.5% BiVO water solution in a quartz cuvette was excited by a 750 nm laser with a 750 nm bandpass filter at various temperatures. The solution was stirred using a magnet at 1400 rpm. The spectra were measured using a spectrometer with a 785 nm longpass filter, placed perpendicularly to the laser beam. Experimentally measured spectra were correlated with temperatures measured by a type K NiCr-Ni thermocouple placed in the water dispersion. Temperatures were changed with steps of 10 °C between 30 and 80 °C. Assuming a linear approximation, the spectra data points can be expanded to form a complete temperature-spectra database for all temperatures between 30 and 80 °C.

The modelling stage includes models for the laser excitation, fiber acceptance region, and temperature distribution. The fluorescent surface is divided into *N* = 640×480 pixels. The spectrum recorded by a fiber with a specific NA, placed in specific *xyz* coordinates is approximated by the following equation:

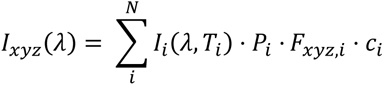

where *I*_*xyz*_(*λ*) is the fluorescence intensity at each wavelength *λ* measured by a fiber in position *xyz*, *i* is the index corresponding to each of the *N* pixels, *I*_*i*_(*λ*, *T*_*i*_) is the fluorescence emitted from the pixel *i* at the temperature *T_i_*, *P_i_* is the laser power intensity, *F*_*xyz*,*i*_ is the fraction of fluorescence emitted by the pixel *i* that is measured by the fiber in position *xyz* (which takes into account acceptance angle of the fiber and distance from source), and *c_i_* is the concentration of nanothermometers in the pixel *i*. All parameters are normalized and range from 0 to 1. The laser power distribution is calculated using the paraxial Helmholtz equation for TEM_00_ Gaussian and top-hat beams.

In the image formation step the algorithm is run multiple times varying the fiber position with fixed laser, temperature and concentration parameters to create the simulated measured image. A 5×5 grid with 500 µm spacing was chosen for the simulation. Moreover, a laser placed in the center of the FOV with beam quality factor *m* = 8, at a distance of 10 mm and beam waist of 1.5 mm was simulated. The nanothermometer concentration was assumed constant and the fiber dimensions were set to NA 0.22 fiber, core diameter 200 µm, distance of 1 mm away from the solder.

In order to produce the training setup, a set of various temperature distributions were created. Skewed Gaussian profiles with different temperature range, position, angle, and width were used due to their similarity to temperature profiles found during laser irradiation. The simulation algorithm was then run for each different temperature profile. The set of values presented in Table 1 was used to create the temperature distributions. The following parameters were varied: the peak position *µ_x_* and *µ_y_*, the two standard deviations *σ_x_* and *σ_y_*, the skewness *a_x_* and *a_y_*, the maximum and minimum temperatures *T*_max_ and *T*_min_, and the rotation *φ*. *T*_min_ was chosen to be randomly assigned in the interval [20, *T*_max_ – 1] with uniform distribution.

**Table 1:** simulation parameters.

| | $\mu_x, \mu_y$ [mm] | $\sigma_x, \sigma_y$ [$10^{-4}$ mm] | $a_x$ | $a_y$ | $T_{\max}$ [$^{\circ}\text{C}$] | $\Phi$ [$^{\circ}$] |
| --- | --- | --- | --- | --- | --- | --- |
| Values | -1 – 1 | 5 – 11 | -2 – 2 | 0 | 40 – 90 | 0 – 90 |
| Step size | 0.5 | 2 | 4 | 0 | 5 | 15 |

The dataset was further improved by discarding samples with low degree of similarity, choosing to use a Structural Similarity Index Measurement (SSIM) of 0.7 as cut-off value. The threshold of 0.7 was carefully selected, representing an optimal balance between preserving data quality and maintaining enough data for training. This approach led to the removal of 10’010 samples out of a total of 38’947, leading to a remaining 28’937 samples, which provide ample data for effective network training.

### CNN architecture and training

The chosen CNN architecture is shown in Figure 3b. Comprised of several convolutional layers, the model integrates upsampling layers between the second and third, and the third and fourth layers. These upsampling layers amplify the image size by a factor of two each time. This action results in an enhancement of resolution, a pivotal step in image upscaling tasks. The activation function selected for this model is the PReLU (Parametric Rectified Linear Unit), a variant of the leaky ReLU equipped with a trainable parameter. This added flexibility enables the PReLU function to adapt its slope during training, allowing the network to discern where non-linearity should be introduced. In terms of loss function, the SSIM (Structural Similarity Index Measure) loss has been favored over the MSE (Mean Squared Error). The motivation behind this choice is the superior performance of SSIM in preserving structures and shapes, which is crucial for our objectives. Serving as the cornerstone of this Convolutional Neural Network (CNN), the convolutional layers are endowed with a set of learnable filters, often referred to as kernels. As the forward pass is carried out, each filter sweeps over the width and height of the input volume, performing a dot product operation between the entries of the filter and the input. This process generates an activation map. Importantly, each layer is tailored to detect different features within the image, with earlier layers often picking up rudimentary features such as edges, while subsequent layers may capture more complex features like textures. During the training phase of the CNN, a configuration of 300 epochs with a batch size of 500 and a learning rate set to the conventional value of 0.001 were used. To optimize the network architecture, the training dataset was divided as follows: a quarter of the entire dataset was reserved for testing, while the remaining 75% was split further into an 80-20 ratio for training and validation, respectively. Following the identification and optimization of the most promising network configuration, it was subjected to an additional round of training involving the complete artificially generated dataset. This comprehensive retraining was executed to enhance the performance of the network when dealing with data derived from laboratory tests.

### CNN experimental validation

A repurposed 3D printer (Ender 2 Pro, Creality) was used as XYZ translation stage with temperature control. A solder paste sample (25% BSA, 6.5% GelMA, 0.3% BiVO, 0.002% TiN, 0.0069% LAP) was placed on the platform of the XYZ translation stage and undergoes heating via laser irradiation. The 750 nm laser, filtered by a 750 nm bandpass filter and focused by a lens, was fixed on the platform and therefore did not move with respect to the paste. The aim of this setup is to mimic the conditions of any multicore fiber, such as one comprising 25 cores, arranged in a quadratic 5×5 configuration. An optical fiber (NA 0.22, ⌀600 µm) interfaced with the spectrometer was placed where the extruder of the 3D printer is usually placed. A 785 nm longpass filter was used to filter the collected radiation.

The measurement phase commences after the sample has been irradiated long enough to reach a stable temperature distribution. A picture with a thermal camera was taken at the beginning and end of the measurement. The stage is used to move the fiber through the 5×5 grid with 500 µm distance between points. The acquisition time of the spectrometer, usually 500 ms, was varied depending on the intensity of the fluorescence, reaching up to 6 s in extreme cases, when the fiber was particularly far from the excited region. The stage bed was also heated to circa 37 °C to simulate the temperature found in the body. The distance between the optical fiber and the sample was kept at around 2 mm. The data was then processed to form the 5×5 low resolution thermal image. The trained CNN was used to reconstruct the thermal distribution and the result was compared to the resized data acquired with the thermal camera. N = 3 for experimental paste samples.

### Thermal imaging through image-guiding fiber

A 54 cm long glass fiber optic image conduit (1534320, SCHOTT) was used to relay the fluorescence image from the solder paste to the NIR cameras. The image conduit has an outer diameter of 1.05 mm and 18k fibers. A medical fiber together with a 750 nm laser were used for excitation. A GRIN (Gradient Index) rod lens (working distance = 0 mm, length = 4.34 mm, diameter = 1.80 mm; #64-525, Edmund Optics) was placed in front of the fiber and moved manually to the position necessary to focus the image. A x10 microscope objective (Bresser) was used to image the other end of the image conduit. A custom 3D-printed filter cube was used to filter out the fluorescence signal from the laser light, as well as to separate the two fluorescence peaks. The following filters were used: 800 nm longpass filter (FELH0800, Thorlabs), 830 nm longpass beamsplitter (F38-830, AHF Analysentechnik), 810 nm bandpass filter (FBH810-10, Thorlabs), 850 nm longpass filter (FELH0850, Thorlabs). For the positioning of the filters see Figure 4a. Two NIR cameras (acA1920-40µm, Basler) were used to record the fluorescence of the two fluorescence peaks. A thermal camera was used as control. A piece of solder paste (25% BSA, 6.5% Gelatin, 0.3% BiVO, 0.002% TiN) on a glass slide was used. The solder paste was placed at a distance between 0.3 and 1 cm away from the GRIN lens. The piece of solder paste was moved manually to simulate the movement of the fiber with respect to the paste during soldering. The laser power was changed between 30 and 80% of the maximum power to verify the performance of the imaging method. The acquisition time was set at 500 ms. The gains were set carefully to avoid saturation and achieve high signal-to-noise ratio. The fluorescence intensity ratios (FIRs) were computed after manually aligning the two images and after removing the background from the images, calculated using the average in a selected area of the images without signal. A Gaussian blur of standard deviation 7 pixels was added to avoid artifacts coming from dark fibers and imperfect image alignment. Moreover, only FIRs from pixels with more than 15 counts in the 810 nm peak image are displayed (all the others are set to 0 by default, appearing dark). The same has been done for thermal images. The calibration was computed through a reference measurement.

The imaging of the thermal distribution was also recorded during soldering of an extender region of the paste. The paste was moved using the *xyz* stage at a speed of 10 mm/min while soldering and imaging were performed. The composite image of the thermal distribution throughout the paste during soldering was created by computing the average temperature at each physical location whenever the temperature at that location was measured and deemed accurate (i.e. in the range 20-80 °C).

### Solder paste recognition ML algorithm

An image segmentation model was trained on a dataset of diverse images of solder pastes of various shapes (up to squares of 1 cm^2^) and nanoparticles concentrations (0.3% BiVO, 0.0002-0.002% TiN) placed on various tissues, such as ex vivo and in vivo porcine blood vessels and intestine. A total of 150 pictures were used, with an 80/20 training/validation split. The images were taken with a smartphone, downscaled to 1920 x 1440 pixels and then labelled manually. The training was performed on a pretrained model (‘yolov8n-seg.pt’ by YOLOv8.0.20, Ultralytics), with 25 epochs, image size of 640 and batch size of 10. More information can be found in Supplementary Information.

### Robotic soldering

A surgical robot (Dexter, Distalmotion SA) controlled through a remote telemanipulator was used. The test was performed inside a laparoscopic trainer simulator box. The optical components used for the temperature-controlled soldering and for the nanothermometry thermal imaging are the ones described in the previous sections. The fibers for the temperature-controlled soldering were integrated with one of the surgical instruments. The image-guiding fiber was inserted using a plastic sleeve through a trocar and moved in position using the robotic arm equipped with a grasper. A metal hook passed through the plastic sleeve aided the movement of the fiber and avoided damage to the fiber from direct grasping. A piece of solder paste was inserted and moved to the desired location using the robotic grasper. A piece of porcine tissue (pig shoulder) was purchased from a local grocery shop and used as model for the experiment. Soldering was then performed and nanothermometry thermal distribution data acquired.

### In vivo soldering

An in vivo study was approved by the Commission of Work with Experimental Animals (project ID: MSMT-15629/2020-4) under the Czech Republic’s Ministry of Agriculture supervision. All procedures were carried out according to Czech as well as regulations of the European Union. A combined intramuscular injection of ketamine (Narkamon 100 mg mL−1, BioVeta a.s., Ivanovice na Hané, Czech Republic) and azaperone (Stresnil 40 mg mL−1, Elanco AH, Prague, Czech Republic) was used to pre-medicate a healthy female Prestice Black-Pied pig (60 months old). Continuous intravenous propofol injection (Propofol 2% MCT/LCT Fresenius Medical Care a.s.) was used to maintain general anesthesia. Nalbuphine (Nalbuphin, Torrex Chiesi CZ s.r.o., Czech Republic) was used intravenously for analgesia assurance. The abdominal cavity was entered following ordinary laparoscopic procedure. The solder paste was cut and placed in a grasper to insert it inside the abdominal cavity. Optical fibers were passed through a 5 mm trocar and moved in position by hand. The intestine and the left lateral abdominal wall (where a defect was made) were soldered using the iSoldering approach, testing pastes of various composition (gelatine- or crosslinked GelMA-based and TiN concentrations of 0.001%, 0.002%, and 0.01%). After euthanasia, tissue samples were collected and chemically fixed in 4% paraformaldehyde in PBS. Histology samples were processed (embedded, sectioned, and stained) by SophistoLab, Muttenz, Switzerland. Samples were imaged using a whole slide scanner at ScopeM, ETH Zurich.

### Mechanical testing

Porcine small intestine was obtained from a local slaughterhouse. The tensile strength was measured using a force meter (AFM-20, TopHomer) and an actuator (PQ12-100-6-R) controlled through a microcontroller (Arduino Uno R4 Minima). Intestine samples were cut into rectangular pieces of 5 ± 2 mm width with a full-width cut placed in the middle. Soldered samples were soldered with a 4 × 4 mm square of GelMA-based solder paste containing a TiN concentration of 0.002%, using a 750 nm laser for up to 10 min at 70 °C. Sutured samples were sutured using a 5-0 polypropylene suture (PROLENE, Ethicon) with one knot. Failure was defined by the full separation of the specimen. Tensile strength was obtained from the breaking load normalized by the width of the specimens. N = 3 for both samples.

### Statistical analysis

No data preprocessing has been employed. Data in bar and line charts are displayed as mean ± SD. The sample size (N) was indicated in the figure captions and Experimental Section. Python has been used for data processing.

## Acknowledgments

We acknowledge funding from the Swiss National Science Foundation (Eccellenza grant no. 181290) and the Oertli Foundation. We thank Dr. Thomas Rduch for his advice and Dr. Vera Kissling for her help with TEM images. We thank Lucas Dosnon for his advice on the image segmentation algorithm. We thank Helen Berti for producing the scientific illustrations (Figure 1a,b). We acknowledge ScopeM, the microscopy center of ETH Zurich, for access to their microscopes.

