## Supplementary information for "Robotic Laser Tissue Soldering for Damage-free Soft Tissue Fusion Guided by Fluorescent Nanothermometry"

**Nanoparticle characterization**

TEM images of BiVO and TiN are see in Figure S1.

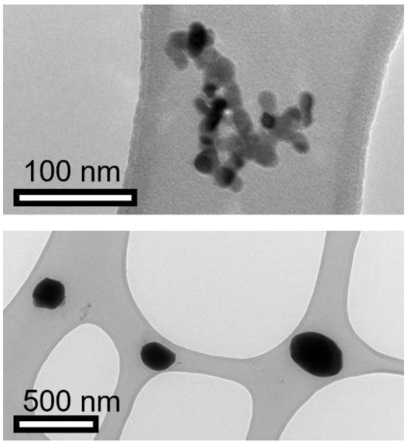


***Figure S1: TEM images.*** *TEM images of TiN (top) and Nd-doped BiVO (bottom).*

### Machine Learning paste detection from image

The training metrics of the machine vision algorithm are shown in Figure S2a. The performance of the algorithm is further demonstrated through additional images used for validation (Figure S2b).

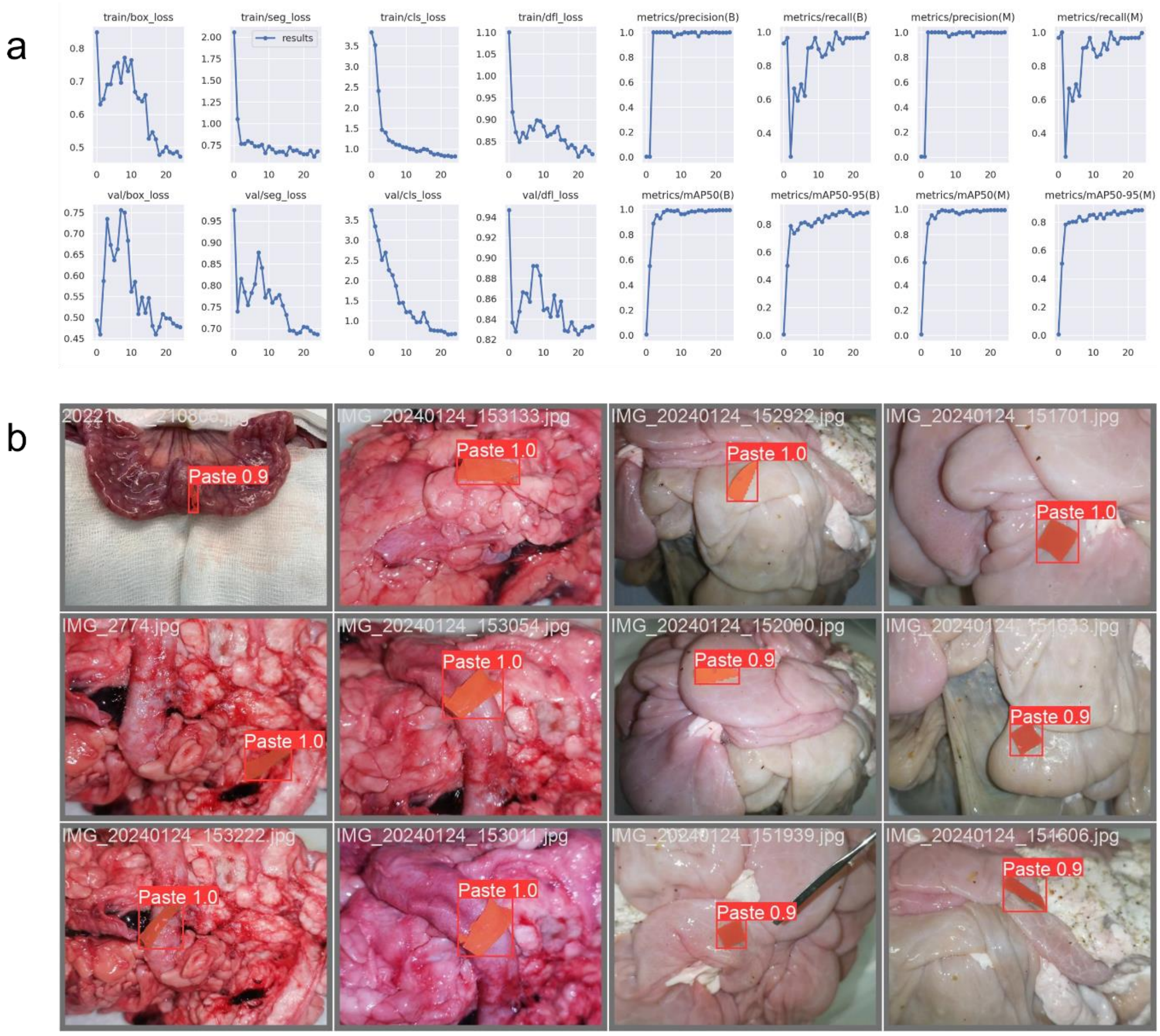

*__Figure S2: machine learning algorithm performance.__ (a) Training and performance metrics of the paste detection and segmentation algorithm. (b) Images from the validation set with the paste being segmented correctly.*

### In vivo minimally invasive surgery with soldering

Additional images of the surgery being performed on the porcine model are shown in Figure S3. It can be noted how the paste can be placed on the desired location (Figure S3a), how the soldering starts when the paste is recognized (Figure S3b), and how the fibers can be moved with graspers to reach areas that are difficult to reach ((Figure S3c), such as the lateral abdominal wall (Figure S3d).

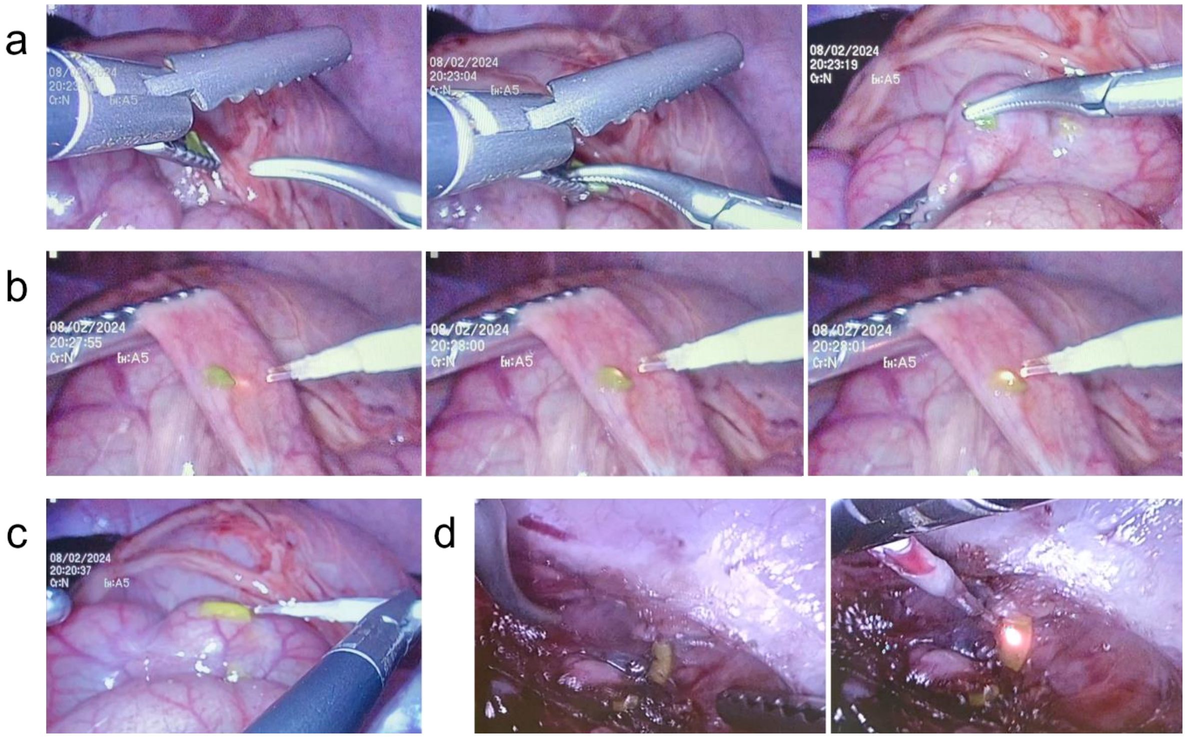


***Figure S3: in vivo laparoscopy images of soldering in chronological order**. (a) Positioning of the paste using the graspers. (b) Soldering of the paste on pig intestine, which starts only after paste detection, as seen from the laser intensity increase on the right image. (c) Fiber positioning using a grasper. (d) Positioning and soldering of paste on the lateral abdominal wall.*
